# Ultrastructure of exospore formation in *Streptomyces* revealed by cryo-electron tomography

**DOI:** 10.1101/2020.07.07.187914

**Authors:** Danielle L. Sexton, Elitza I. Tocheva

**Affiliations:** Department of Microbiology & Immunology, Life Sciences Institute, The University of British Columbia, Vancouver, BC, Canada

**Keywords:** microbial ultrastructure, cryo-electron tomography, Streptomyces, multicellular bacteria, filamentous bacteria, sporulation, cell envelope, bacterial cytoskeleton

## Abstract

Many bacteria form spores in response to adverse environmental conditions. Several sporulation pathways have evolved independently and occur through distinctive mechanisms. Here, using cryo-electron tomography (cryo-ET), we examine all stages of growth and exospore formation in the model organism *Streptomyces albus*. Our data reveal the native ultrastructure of vegetative hyphae, including the polarisome and filaments of ParA or FilP. In addition, we observed septal junctions in vegetative septa, likely involved in effector translocation between neighbouring cells. During sporulation, the cell envelope undergoes dramatic remodeling, including the formation of a spore cortex and two protective proteinaceous layers. Mature spores reveal the presence of a continuous spore coat and an irregular rodlet sheet. Together, these results provide an unprecedented examination of the ultrastructure in *Streptomyces* and further our understanding of the structural complexity of exospore formation.

## Introduction

Sporulation is a developmental process that culminates in the production of specialized dormant life forms called spores, which are readily dispersed in the environment. Spores are morphologically distinct from vegetative cells, often having additional protective structures on the surface such as modified peptidoglycan (PG) and several proteinaceous layers. The decreased water content and low metabolic activity of spores further enhance their resistance to environmental stressors (Constant et al., 2008;Elliot and Flardh, 2012;Ghosh et al., 2015;Liot and Constant, 2016). Several independent mechanisms for bacterial sporulation have evolved. The most extensively characterized mode of sporulation, both genetically and structurally, is endospore formation in Firmicutes, exemplified by *Bacillus subtilis* (Higgins and Dworkin, 2012;Tocheva et al., 2013;Khanna et al., 2019). Exospore formation, on the other hand, has been extensively characterized in the multicellular bacterium *Streptomyces*, member of the phylum Actinobacteria. *Streptomyces* grow vegetatively as a series of interconnected multinucleate compartments, forming multicellular branching filamentous hyphae. Nutrient limitation triggers sporulation and the process begins by the emergence of specialized non-branching aerial hyphae from the colony surface (McCormick and Flardh, 2012) (Fig. 1). The hyphae undergo synchronous cell division to produce numerous identical spores. Mature spores are released into the environment to ensure dispersal of genomic material. While *Streptomyces* sporulation is phenotypically similar to many filamentous fungi, these processes are likely the result of convergent evolution.

**Figure 1.**
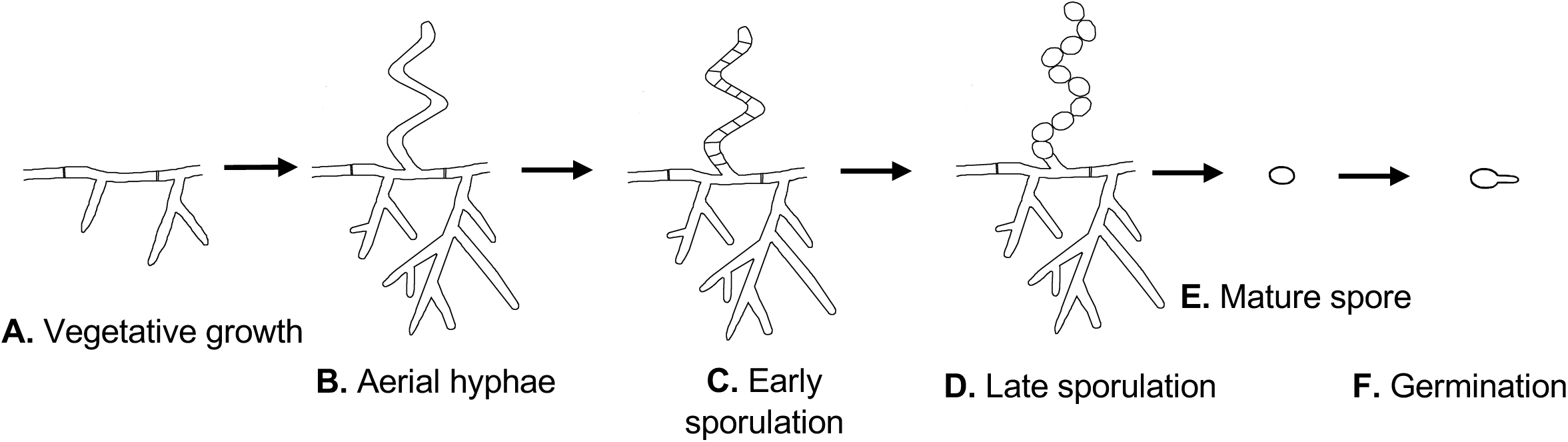
*Streptomyces* life cycle. Schematic representation of the major growth stages: (A) vegetative hyphae, (B) early sporulation begins with aerial hyphae formation, (C) synchronous septa formation for immature spores, (D) spore maturation, (E) release of mature spores, and (F) germination into vegetative hyphae.

In the soil, *Streptomyces* exists predominantly as spores that remain dormant until favourable growth conditions are sensed (Ensign, 1978). The exact nutrient set required for germination remains unknown, however some studies have shown that divalent cations such as Ca^2+^ and Mg^2+^ can induce germination (Ensign, 1978;Eaton and Ensign, 1980). Following the cue to initiate germination, spores rehydrate and switch from phase-bright to phase-dark when viewed with phase-contrast light microscopy. This stage proceeds without new cell wall synthesis (Ensign, 1978). When a new vegetative cell emerges from the spore in the form of a germ tube (Fig. 1F), degradation of the spore cortex and synthesis of new PG at the tip is required (Flardh, 2003;Sexton et al., 2020). Polar growth of *Streptomyces* is directed by the polarisome, a complex containing DivIVA, Scy, and FilP proteins (Flardh, 2003;Fuchino et al., 2013;Holmes et al., 2013) (Fig. 1A). DivIVA is a coiled coil protein that forms filaments at sites of negative curvature (Hempel et al., 2008;Lenarcic et al., 2009) and interacts directly with the inner leaflet of the cytoplasmic membrane (Stahlberg et al., 2004;Oliva et al., 2010). Scy is another coiled coil protein that forms filaments, and its interaction with DivIVA is thought to control the frequency of branching during vegetative growth (Holmes et al., 2013) as well as the anchoring of the chromosome to the hyphal tip (Ditkowski et al., 2013;Kois-Ostrowska et al., 2016). The last member of the polarisome, FilP, is shown to form intermediate filaments and aid in several important molecular processes, including DivIVA stabilization (Frojd and Flardh, 2019) and hyphal tip rigidity (Bagchi et al., 2008;Fuchino et al., 2013). Furthermore, the polarisome guides the localization of penicillin binding proteins and the glycosyltransferase CslA involved in the synthesis of new PG and cellulose-like polymer, respectively (Xu et al., 2008;Holmes et al., 2013;Ultee et al., 2020). Overall, the polarisome is a key protein complex that directs tip extension, localizes chromosomes, produces extracellular polysaccharides, and organizes the cytoskeleton.

Filamentous bacteria, such as cyanobacteria, have evolved septal junctions that facilitate communication between neighbouring cells (Merino-Puerto et al., 2011;Wilk et al., 2011). Similarly, in *Streptomyces*, vegetative hyphae are occasionally subdivided into linked syncytial compartments by vegetative septa (McCormick and Flardh, 2012). Vegetative septa are semi permeable, as plasmid DNA and GFP have been shown to traverse vegetative septa (Kataoka et al., 1991;Hopwood and Kieser, 1993;Celler et al., 2016). While vegetative septa are not required for growth or viability (McCormick et al., 1994), they may confer other advantages. For example, as the colony prepares for sporulation, vegetative hyphae in the centre of the colony undergo lysis (Miguelez et al., 1999). This is thought to provide valuable nutrients that fuel sporulation. Compartmentalizing the vegetative hyphae with selectively-permeable septa could allow for secure transport of valuable nutrients through the cytoplasm of the hyphal network to sites of sporulation. Notably, the presence of septal junctions in *Streptomyces* vegetative septa that could allow for transport of molecules has been suggested but not confirmed.

The first step in sporulation is the production of aerial hyphae above the colony surface (Fig. 1B). In order to lower the surface tension at the air-colony interface, the aerial hyphae are coated with a hydrophobic layer composed of rodlin and chaplin proteins (Claessen et al., 2003;Elliot et al., 2003;Claessen et al., 2004). Without this rodlet layer, colonies are unable to produce aerial hyphae or sporulate (Claessen et al., 2003;Elliot et al., 2003). Similar to vegetative growth, upwards extension is presumably directed by DivIVA (Flardh and Buttner, 2009). Once aerial hyphae cease lengthening, they are subdivided by septa to produce immature spores (Fig. 1C). Septal formation is directed by FtsZ, which synchronously forms numerous Z rings 1-2 μm apart inside the aerial hyphae (McCormick et al., 1994). MreB localizes to the formed septa, possibly to aid in PG synthesis at the newly-formed septa (Mazza et al., 2006). Subsequently, MreB and other components of the *Streptomyces* spore-wall synthesizing complex (SSSC) localize around the entire spore to direct cortex synthesis and spore maturation (Fig. 1D)(Mazza et al., 2006;Kleinschnitz et al., 2011). It is yet unclear how much of the existing PG is modified to become a part of the spore cortex. Similar to endospores, it is hypothesized that a proteinaceous coat is formed during spore maturation on the surface of *Streptomyces* spores, however, it has not been observed to date. Once the maturation process is complete, spores are released from the spore chain, likely by mechanical forces (Fig. 1E).

Previous characterization of *Streptomyces* development has been done using traditional electron microscopy (EM) techniques. These techniques rely on dehydration and crosslinking by fixation, which can disrupt cellular ultrastructure. With this study, we characterized all stages of *Streptomyces* growth and development using cryo-electron tomography (cryo-ET). Cryo-ET preserves cells in their native state and provides three dimensional reconstructions of whole cells at ∼4 nm resolution (Tocheva et al., 2010). By cryogenically preserving *S. albus* at different stages of their life cycle, we aimed to characterize the ultrastructure associated with vegetative and sporulating cells. Our observations include the structure of the polarisome in vegetative hyphae, septal junctions between vegetative cells, the rodlet layer on the surface of aerial hyphae and spore chains, and the presence of a spore coat surrounding mature spores. Therefore, cryo-ET provides a unique opportunity to directly observe notable cellular structures in *Streptomyces* that will ultimately initiate new lines of research.

## Materials and Methods

### Strains and Growth Conditions

*S. albus* was grown on solid soy flour mannitol agar (20 g/L soy flour, 20 g/L mannitol, and 20 g/L agar) or in liquid 2 x YT medium (16 g/L tryptone, 10 g/L yeast extract, 5 g/L sodium chloride) supplemented with 20 mM MgCl2 (Hopwood et al., 2000).

### Sample preparation

For vegetative and sporulation samples, cells were grown on soy flour mannitol agar until the desired growth stage was reached. For germination, spores were resuspended in liquid 2 x YT medium supplemented with 20 mM MgCl2. Spores were heat shocked at 50 °C for 10 min and then incubated at 30 °C for 4 hours, until germ tubes were observed with light microscopy. Prior to imaging, single colonies were picked off the plate and resuspended in phosphate buffered saline pH 7.2. Images were collected using phase contrast microscopy on an upright Ziess Axio Examiner Z1 equipped with an Axiocam 506 mono at 1000x total magnification and processed using Zen Blue 2.1. To measure cell compartment lengths, membranes were stained with 1/1000 dilution of CellBrite Fix 640 (Biotium). Cells were measured using Fiji software (Schindelin et al., 2012).

### Cryo-ET data collection and processing

Samples were mixed with 20-nm colloidal gold particles, loaded onto glow-discharged carbon grids (R2/2, Quantifoil) and plunge-frozen into liquid ethane-propane mix cooled at liquid nitrogen temperatures with a Mark IV Vitrobot maintained at room temperature and 70% humidity. Tilt series of samples at all growth stages were collected using SerialEM (Mastronarde, 2005) on a Titan Krios 300 keV transmission electron microscope (Thermo Fischer Scientific) equipped with a Falcon III camera, Gatan K3 camera, or a Gatan K3 camera and Bioquantum energy filter. Tilt series were collected at 10 ⍰m defocus, 120 e^-^/Å^2^ total dose, ⍰ 60° tilt, and 1° increments. Three dimensional reconstructions were calculated using the IMOD package and the back-weighted projection method (Kremer et al., 1996).

## Results and Discussion

To characterize exospore formation in Actinobacteria, we collected tomograms from each stage of *Streptomyces* growth (Fig. 2). Model *Streptomyces* species such as *S. coelicolor* and *S. venezuelae* produce cells that are too thick (∼ 1 ⍰m) for direct imaging with cryo-ET, so we used *S. albus* as a model system. *S. albus* vegetative hyphae are <0.5 ⍰m in diameter and thus suitable for analysis by cryo-ET without additional thinning of the sample.

**Figure 2.**
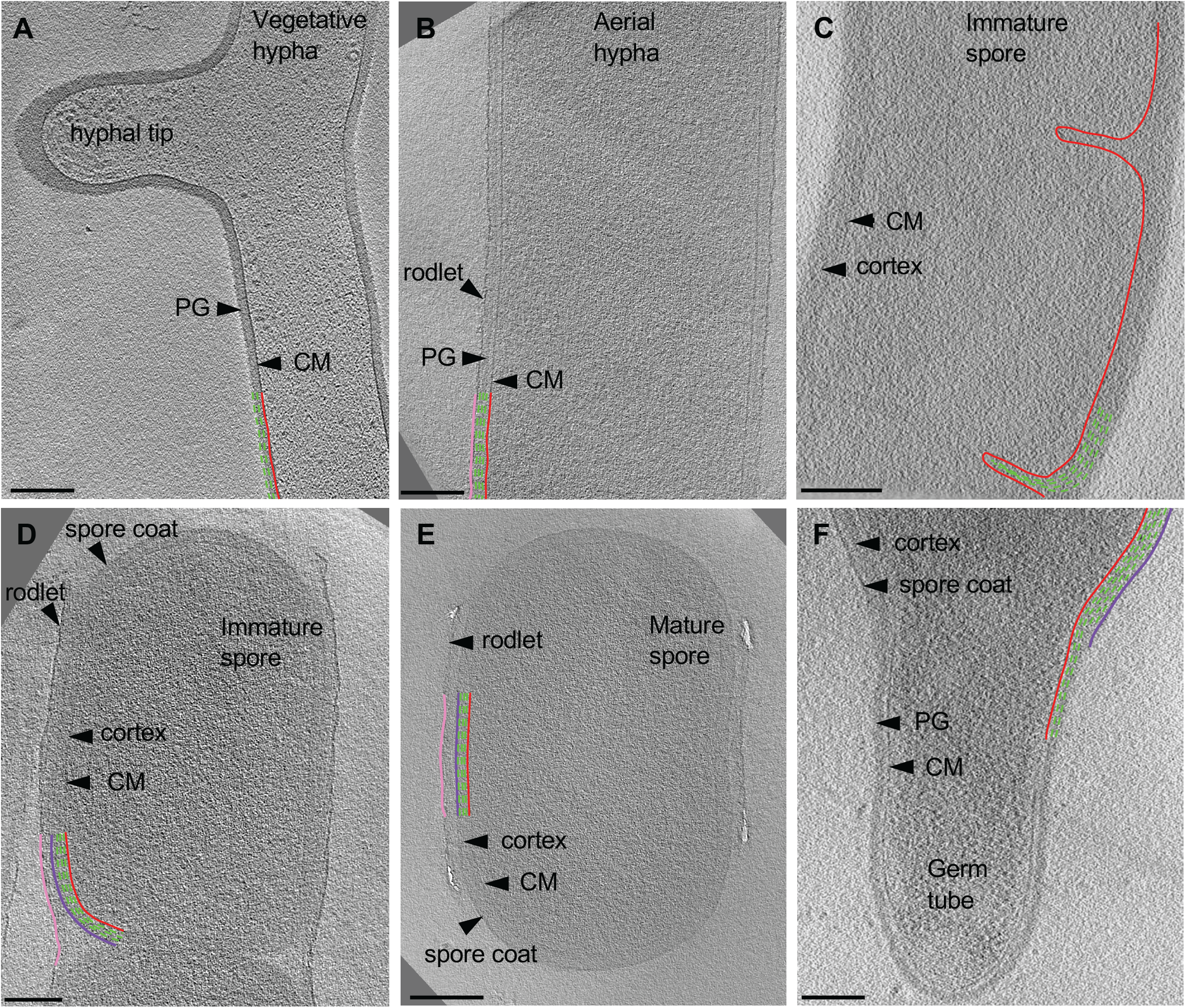
Cryo-ET of vegetative, sporulating and germinating *Streptomyces*. Tomographic slices through S. albus at different growth stages corresponding to (A) vegetative hyphae, (B) aerial hyphae formation during early sporulation, (C) septa formation during sporulation, (D) spore maturation, (E) release of mature spores, and (F) germination. Cytoplasmic membrane (CM) is shown in red. Vegetative peptidoglycan (PG) and cortex are shown in green. Rodlet ultrastructure and spore coat are shown in pink and dark purple, respectively. Each tomographic slice is 20-nm thick. Scale 200 nm.

### Features of vegetative hyphae

First, we sought to characterize vegetative growth, and subsequently identify major cellular changes induced by sporulation. Vegetative hyphae were ∼ 0.5 ⍰m in diameter and ran up to tens of microns in length, with occasional cross walls and membranes separating the hyphae into 10 μm long multinucleate compartments (Table 1). The vegetative PG was ∼35 nm thick (Table 1, Fig. 4) with an exterior surface that appeared smooth and undecorated (Fig. 2A). At hyphal tips, the PG was 5-20 nm thicker than the lateral region of the same tip, presumably due to deposition of the cellulose cap to protect the nascent PG during synthesis (Fig. 2A, Table 1). Other than the variation of thickness at the tip, the cell wall appeared uniform (Fig. 2A). Our results were consistent with reports on *S. coelicolor* sacculi, where the apical region appeared thicker than the lateral wall (Ultee et al., 2020); however, we did not observe distinct lamellae of cellulose-like polymers and PG at the hyphal tips in *S. albus*. This difference in observations could be due to the experimental conditions used by the two studies. While both studies used cryo-ET, we visualized whole cells under turgor pressure whereas Ultee *et al* imaged thin purified sacculi that were boiled in SDS prior to imaging.

**Table 1.**
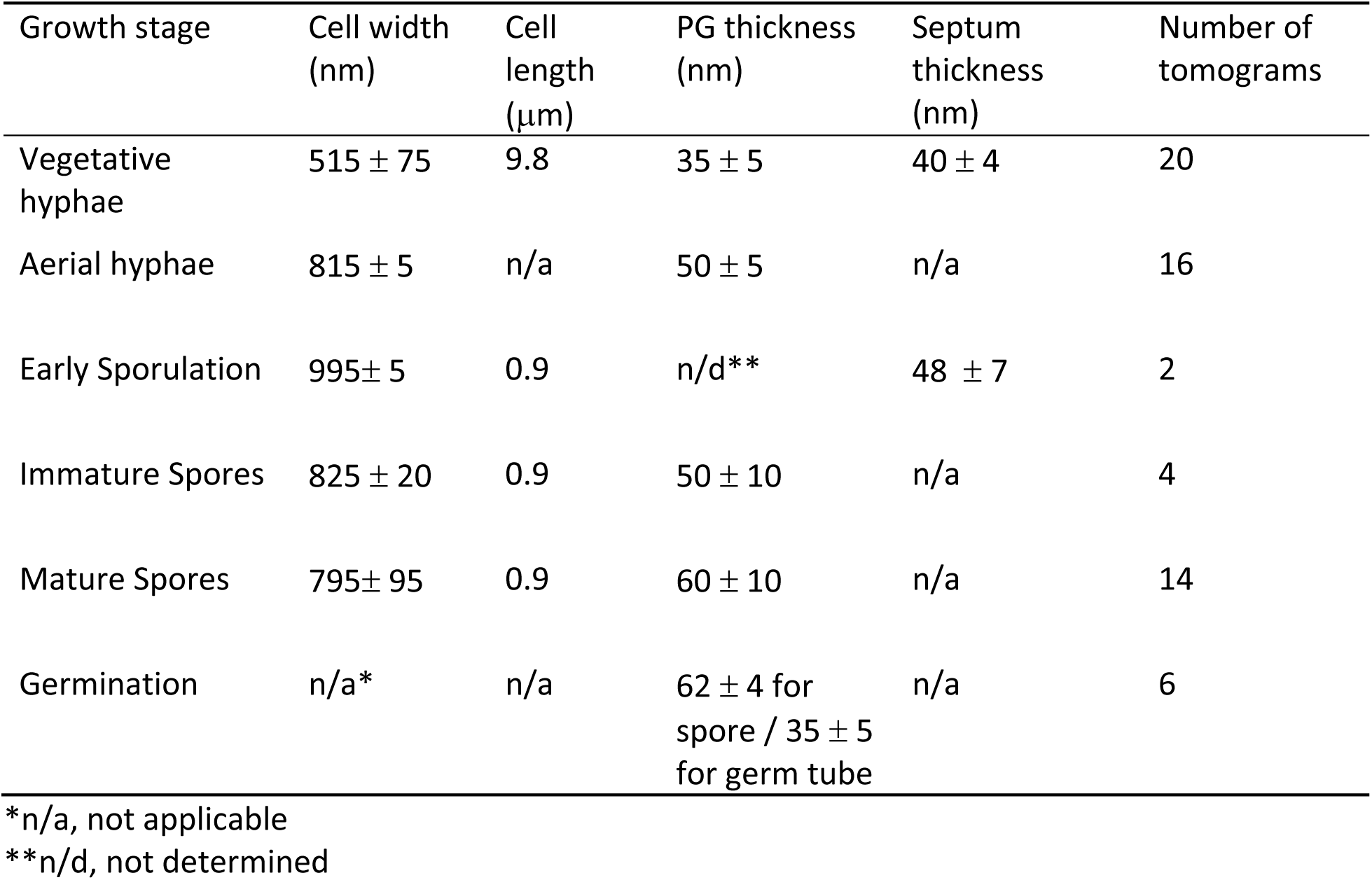
Cell dimensions, PG and cortex thickness and septum thickness at each developmental stage.

| Growth stage | Cell width (nm) | Cell length (μm) | PG thickness (nm) | Septum thickness (nm) | Number of tomograms |
| --- | --- | --- | --- | --- | --- |
| Vegetative hyphae | 515 ± 75 | 9.8 | 35 ± 5 | 40 ± 4 | 20 |
| Aerial hyphae | 815 ± 5 | n/a | 50 ± 5 | n/a | 16 |
| Early Sporulation | 995 ± 5 | 0.9 | n/d** | 48 ± 7 | 2 |
| Immature Spores | 825 ± 20 | 0.9 | 50 ± 10 | n/a | 4 |
| Mature Spores | 795 ± 95 | 0.9 | 60 ± 10 | n/a | 14 |
| Germination | n/a* | n/a | 62 ± 4 for spore / 35 ± 5 for germ tube | n/a | 6 |

\*n/a, not applicable
\*\*n/d, not determined

**Figure 3.**
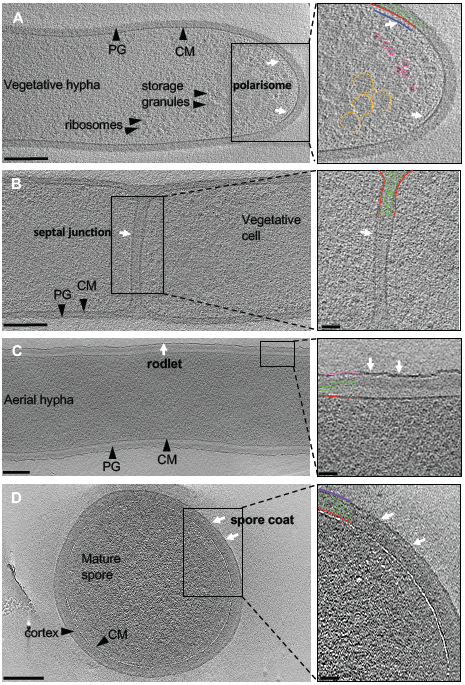
Ultrastructure of *Streptomyces*. A) A tomographic slice through a hyphal tip. The cytoplasmic membrane (red) and peptidoglycan (green) form the cell envelope. Features: polarisome (blue), glycogen storage granules (yellow), and ribosomes (pink). (B) Septal junction connecting two neighboring cells (white arrow). (C) The rodlet layer (pink) on the surface of aerial hyphae and immature spores. (D) A 10-nm thick spore coat (purple) is visible on the surface of mature spores. Insets show a magnified image of the boxed areas. Scale bar 200 nm, inset 50 nm.

**Figure 4.**
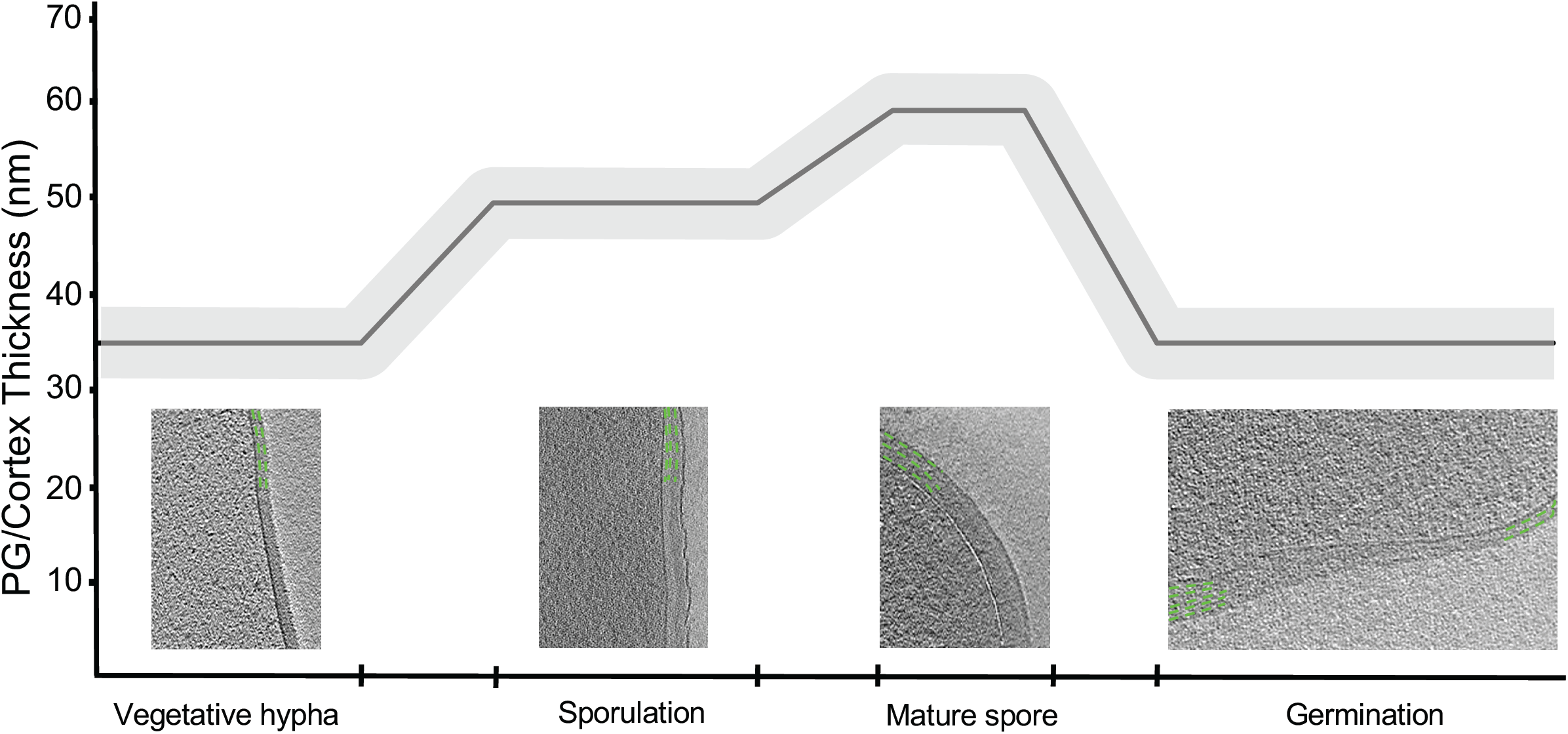
PG and cortex remodeling during exospore formation. During sporulation, PG thickness increases from ∼35 nm in vegetative cells to ∼50 nm in aerial hyphae. During spore maturation, the cortex is thickened to ∼60 nm. During germination, the vegetative PG is a continuation of the inner layers of the spore cortex. PG and spore cortex are shown in green. Each tomographic slice is 20-nm thick.

Ribosomes and storage granules were observed throughout vegetative hyphae (Fig, 3A). The storage granules in *S. albus* were likely composed of glycogen as glycogen has been shown to accumulate during vegetative growth to fuel sporulation (Brana et al., 1986;Rueda et al., 2001). We observed a 6-nm wide filament ∼10 nm underneath the cytoplasmic membrane at hyphal tips that was not detected elsewhere in the vegetative hyphae (Fig. 3A, inset). We speculate that the layer represents the polarisome and is composed of DivIVA, Scy and FilP. While purified FilP formed striated bundles *in vitro* when imaged with negative staining EM (Javadi et al., 2019), we did not observe such structures in our tomograms, suggesting that FilP may adopt alternate conformations under native conditions or form structures below the resolution limit of the technique. Additional filaments, likely ParA (Kois-Ostrowska et al., 2016), were occasionally observed in the cytoplasm of the vegetative hyphae near hyphal tips. Collectively, our observations highlight the complex interplay of cytoskeletal proteins at the hyphal tip to co-ordinate growth and chromosome positioning inside the cell.

*Streptomyces* hyphae are subdivided into compartments by occasional septa and cross membranes. Cross membranes were observed in vegetative hyphae and described previously (Celler et al., 2016;Yague et al., 2016). Our tomograms showed that vegetative septa divided hyphae into compartments without completing cell division (Fig. 3B). PG in septa was continuous with the vegetative cell wall and appeared visually similar, suggesting comparable composition and structure of the two. The septa appeared slightly thicker (∼50 nm) than the surrounding cell wall of ∼35 nm (Table 1), which could be due to the difference in the macromolecular machinery guiding the processes (McCormick et al., 1994;Mistry et al., 2008). Several cryotomograms of vegetative septa revealed 12-nm wide septal junctions with a 9-nm wide lumen (Fig. 3B inset), which could allow free flow of small molecules, proteins, and DNA between adjacent cells. Structures of similar septal junctions composed by Fra family proteins and SepJ were recently reported in filamentous cyanobacteria (Weiss et al., 2019). Homologs of these proteins do not exist in *S. albus*, suggestive of a different mechanism for cell-to-cell communication in *Streptomyces*.

### The cell envelope undergoes dramatic remodeling during sporulation

Sporulation in *Streptomyces* is triggered by nutrient limitation (McCormick and Flardh, 2012). The first step of the process is directed by the polarisome and involves the formation of morphologically distinct aerial hyphae. The aerial hyphae of *S. albus* were on average 300-nm thicker than the vegetative cells (Table 1). As a result, less detail was resolved in the cytoplasm of these cells. Some of the notable changes were observed in the cell envelope morphology and composition. The aerial hyphae had thicker PG compared of vegetative cells (Fig. 2B and C, Fig. 4, Table 1). In addition, the rodlet layer that is integral to the formation of sporulating cells (Claessen et al., 2002;Claessen et al., 2003;Elliot et al., 2003) was observed on the cell surface of aerial hyphae (Fig. 3C). During aerial hyphae growth, the rodlet layer appeared discontinuous (Fig. 3C, inset), reflective of the overlapping basket weave structure previously reported on the surface of aerial hyphae and mature spores (Claessen et al., 2003;Elliot et al., 2003).

Cryotomograms of aerial hypha showed synchronous formation of sporulative septa and the development of immature spores (Fig. 2C). At this stage, the cells were ∼1 ⍰m in diameter, ∼200-nm thicker than aerial hyphae. Our data showed ∼50-nm thick sporulative septa ∼1 ⍰m apart but, due to the thickness of these cells, we were unable to clearly resolve the rodlet layer and PG on the cell surface. (Table 1, Fig. 2C). Following septation, aerial hyphae divide into immature spores via a mechanism described as ‘V snapping’, which relies on turgor pressure and structural weakening of the PG to drive cell division in milliseconds (Zhou et al., 2016). In immature spore chains, the rodlet layer remained intact as a 20-nm thick sheath (Fig. 2D). At this stage, an additional, ∼10-nm thick electron dense layer, likely the spore coat, was observed surrounding each spore between the cortex and the rodlet layer (Fig. 3D). The identification of the proteins comprising the spore coat is of significant interest, as it would allow for the characterization of the role of this layer in spore survival.

Spore maturation is defined by extensive remodeling of the cortex, condensation of the chromosome, deposition of spore pigments, and the entrance into dormancy (Bobek et al., 2017). Once matured, spores disperse from the spore chain presumably via mechanical forces. In our cryotomograms, the spore cortex appeared slightly thicker, increasing from ∼50 nm to ∼60 nm, during maturation (Table 1, Fig. 2D and E, Fig. 4), likely directed by the SSSC (Kleinschnitz et al., 2011). In some instances, two distinct cortex layers could be identified - a darker inner cortex (∼20-nm thick) and a lighter, outer cortex (∼30-nm thick). The spore cortex overall has an increased percentage of 3-4 crosslinks and decreased 3-3 crosslinks compared to the PG of aerial hyphae (van der Aart et al., 2018), however, the structural differences between the inner and outer cortices remain unknown.

There has been speculation about whether the rodlet layer functions as a spore coat. For a spore coat to be protective, it needs to completely surround the spore and remain tightly associated with it. Our cryotomograms showed that after dispersal, the rodlet layer sporadically remained associated with the sides of some spores (Fig. 2E), while the novel spore coat remained intact even on spores where the rodlet layer appeared detached (Fig. 3D). The rodlet layer may have additional functions following spore dispersal, including promoting interactions with the flagella of motile soil bacteria (Muok et al., 2020).

### Germination

Conditions that prompt spore germination are still undefined, however, heat shock and divalent cations such as Ca^2+^ and Mg^2+^ are known to stimulate germination (Hirsch and Ensign, 1976;Eaton and Ensign, 1980). Since germination defines the transition between sporulation and vegetative growth, it allowed us to make direct comparisons between the two stages. In cells undergoing germination, the cortex was ∼60 nm thick whereas the PG of the germ tube appeared ∼ 35 nm thick (Fig. 4, Table 1). These measurements are comparable to those for mature spores and vegetative hyphae, respectively. The vegetative PG appeared as a continuation of the inner layers of the cortex, and the outer layers of the cortex and spore coat could be seen peeling back from where the germ tube emerged (Fig. 2F). This observation suggested that the inner and outer cortices have different structures and roles during the sporulation and germination processes.

## Conclusions

Our study shows the complete life cycle of *Streptomyces* from vegetative growth, through sporulation and germination in unprecedented detail revealed by cryo-ET. We identified septal junctions between vegetative cells, the polarisome directing polar growth, macromolecular changes involving membrane movement, PG and cortex synthesis, as well as the ultrastructure of the spore coat and rodlet layer.

## Author Contributions

**DS** and **ET** contributed equally to the manuscript. **DS** and **ET** designed experiments, performed work, processed cryo-ET data, revised the manuscript. **DS** wrote the first draft of the manuscript. **ET** coordinated the project.

## Acknowledgements

We would like to thank Dr. Claire Atkinson and the High Resolution Macromolecular Cryo-Electron Microscopy facility at the University of British Columbia, Dr. Mike Strauss and the Facility for Electron Microscopy Research at McGill University, and Dr. Craig Yoshioka at the Pacific Northwest Cryo-EM Center for assistance with microscope operation and data collection.

